# Identifying Antisense Oligonucleotides for Targeted Inhibition of Insulin Receptor Isoform A

**DOI:** 10.1101/2025.03.06.641707

**Authors:** Christopher A. Galifi, Ryan J. Dikdan, Divyangi Kantak, Joseph J. Bulatowicz, Krystopher Maingrette, Samuel I. Gunderson, Teresa L. Wood

**Affiliations:** Department of Pharmacology, Physiology, & Neuroscience, Center for Cell Signaling and Cancer Institute of New Jersey, Rutgers Biomedical and Health Sciences, Newark, NJ, United States; Public Health Research Institute, Rutgers Biomedical and Health Sciences, Newark, NJ, United States; Department of Molecular Biology and Biochemistry, Nelson Biological Laboratories, Rutgers University, Piscataway, NJ, United States

**Keywords:** insulin receptor, IR-A, antisense oligonucleotides, breast cancer, cancer therapeutics

## Abstract

**INTRODUCTION:** The insulin receptor (IR) is alternatively spliced into two isoforms, IR-A and IR-B. IR-B is primarily associated with metabolic signaling, whereas IR-A is highly expressed during embryogenesis. IR-A specifically has been associated with several aggressive cancers; however, selective targeting of IR-A has proven difficult due to its homology with IR-B.

**METHODS:** We generated several antisense oligonucleotides (ASOs) that target the exon 10-12 splice junction site present in IR-A, but not IR-B, mRNA. To test the efficacy of the ASOs, we performed lipofectamine transfections of MDA-MB-231 breast cancer, 22Rv1 prostate carcinoma, and Hs822.T Ewing sarcoma cell lines. We also incubated the MDA-MB-231 cell line with the ASOs in the absence of lipofectamine to determine if they are taken into cells unassisted.

**RESULTS:** One ASO variant selectively reduced IR-A mRNA levels with minimal impact on IR-B mRNA and significantly reduced total IR protein. The IR-A ASO successfully induced selective IR-A knockdown in MDA-MB-231 breast cancer cells, which was maintained after a one-week incubation with the ASO. The ASO selectively reduced IR-A mRNA when administered to cells in high doses without the use of a vehicle (i.e. gymnotic delivery). The ASO was also effective at reducing IR-A mRNA in Hs822.T Ewing Sarcoma and 22Rv1 prostate carcinoma cells.

**DISCUSSION:** We have developed an ASO that targets IR-A with minimal off-target knockdown of IR-B. We hypothesize that the IR-A ASO will be a useful research tool and may have therapeutic value by inhibiting the oncogenic functions of IR-A in cancer cells.

## Introduction

The insulin receptor (IR) is a receptor tyrosine kinase that primarily stimulates glucose metabolism, among other cellular processes (1). IR expression is correlated with worse breast cancer prognosis, and as such, is a viable target for cancer treatment (2–4). Breast cancer is the most common cancer in women, and its incidence and disease burden continue to rise (5). When breast cancer patients present with distant metastasis at diagnosis, their 5-year relative survival rate is only 32.4% (6).

The IR consists of an extracellular α subunit and an intracellular and transmembrane β subunit; the primary RNA transcript undergoes alternative splicing into the A (IR-A) or B (IR-B) isoforms that differ by one exon (exon 11 encoding 12 amino acids of the α subunit) (7). For a detailed structural illustration of the IR protein, please refer to Figure 1 and 2 of the review by Belfiore and colleagues (8). IR-A, commonly expressed in fetal and cancer cells, stimulates the proliferation of both epithelial and mesenchymal cells through insulin-like growth factor 2 (IGF2) binding (9). The IR-B isoform, which is highly expressed in metabolic tissues, is the primary regulator of glucose homeostasis. IR-A has a greater affinity for IGF2 than does IR-B; the IR-A-IGF2 signaling loop is used by several cancer types to promote mitogenesis (10, 11). A high IR-A to IR-B ratio is common in breast cancer cell lines and in human breast tumors at all stages (3, 7, 12–15), and high IR-A expression is associated with worse cancer prognosis in a number of cancer types including breast, prostate, and endometrial cancer (8). In LCC6 breast cancer cells, nonselective knockdown of both IR isoforms via shRNA alleviated tumorigenic hallmarks *in vitro,* as well as metastasis in a mouse xenograft model (16). Inhibiting total IR systemically causes insulin resistance due to IR-B metabolic functions; thus, developing IR-A specific research tools and inhibitors will permit further study of its function in cancer and introduce potential cancer therapeutics (4).

**Figure 1.**
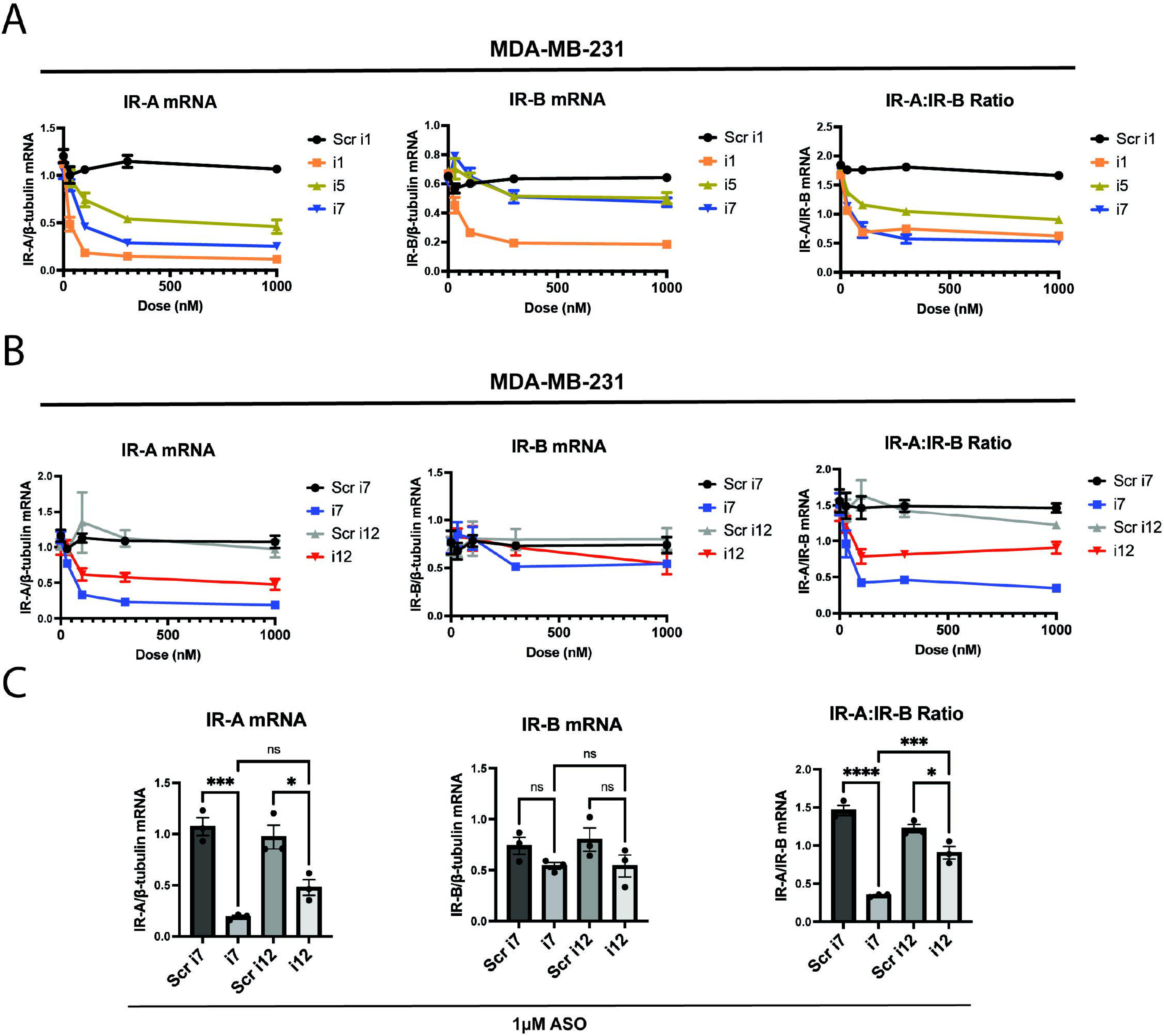
The anti-IR-A ASO “i7” reduces IR-A mRNA selectively without significantly reducing IR-B mRNA up to 1 µM. ASOs were transfected via Lipofectamine RNAiMAX reagent and cells were isolated for processing 24 hours post-transfection. (A) Dose-response and qPCR analysis of a 5-10-5 gapmer (i1), 4-10-4 gapmer (i7), and 5-8-5 gapmer (i5) targeting the exon 10-12 junction of IR-A in MDA-MB-231 triple-negative breast cancer cells. IR-A mRNA (left), IR-B mRNA (middle), and the IR-A:IR-B mRNA ratio (right) were assessed using primers as previously described (Flannery et al., 2016) relative to β-tubulin levels. (B) i7 was compared to a scramble-sequence control and an LNA-modified ASO, i12. IR isoform expression was assessed as in (A). (C) Bar plots of 1 µM i7 and i12 transfections were compared via one-way ANOVA, assessing changes in IR-A mRNA (left), IR-B mRNA (middle), and the IR-A:IR-B mRNA ratio (right).

**Figure 2.**
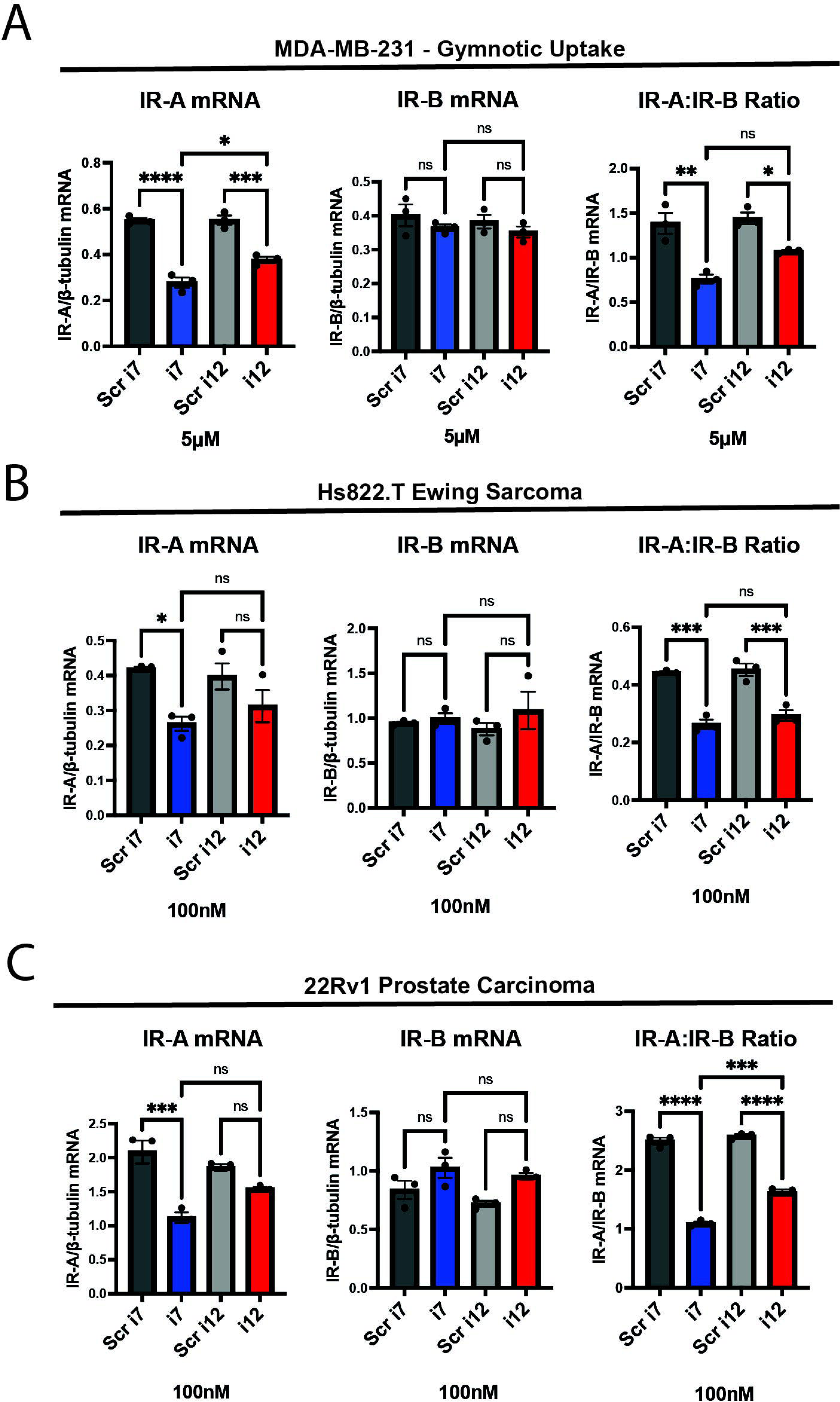
i7 can be taken into cells via gymnosis without a delivery vehicle and selectively knocks down IR-A in other cancer cell types. (A) MDA-MB-231 cells were incubated with 5 µM of i7 and i12, without lipofectamine, for 24 hours before harvest and qPCR analysis of IR-A mRNA (left), IR-B mRNA (middle), and the IR-A:IR-B mRNA ratio (right). (B) Hs822.T Ewing sarcoma cells and (C) 22Rv1 prostate carcinoma cells were transfected with 100 nM i7 and i12 via lipofectamine delivery. IR-A (left), IR-B (middle), and IR-A:IR-B (right) mRNA were assessed via qPCR as previously described.

Inhibiting IR-A specifically poses a challenge due to its high homology to IR-B at the mRNA and protein level. Currently, there is no established method to selectively target IR-A at the protein level, and there is no antibody available that selectively binds IR-A versus IR-B. Splice-switching oligonucleotides (SSOs) have been developed that modulate the IR-A:IR-B ratio (17); however, while these oligonucleotides permit more rigorous study of the unique functions of each IR isoform, the CUG-BP1-blocking SSO that decreases IR-A mRNA also causes a reciprocal increase in IR-B mRNA. For comprehensive description and illustration of the mechanism of action of SSOs, please refer to the review by Havens and Hastings (18). While inhibiting IR-A function with SSOs may have therapeutic value in certain cancers, increasing IR-B metabolic signaling could be detrimental, as IR-B mRNA expression also correlates with worse breast cancer outcomes (3). Thus, developing a reagent that can selectively target IR-A with minimal impact on IR-B expression has potential as both a research tool and therapeutic agent.

Distinguishing IR-A and IR-B protein levels remains a major gap in the IR field; however, primers have been developed that can reliably determine IR-A versus IR-B mRNA expression (19, 20). For a detailed illustration of IR-A and IR-B mRNA, the qPCR primer assay for IR-A and IR-B mRNA quantification, as well as the Exon 10-12 junction that we used for IR-A ASO targeting, please refer to Figure 1 in the article by Flannery and colleagues (19). In a previous study designed to knock down IR-A and IR-B mRNA using siRNAs, the investigators reported that only the anti-IR-B siRNA was successful (21). As such, our goal was to selectively knock down IR-A mRNA using an extensively modified ASO targeting the IR exon 10-12 junction. ASOs silence genes via RNase H-mediated mRNA degradation and steric blockage of translation (22). A complete review and visualization of these mechanisms can be found in the article by Crooke (23). A wide variety of chemical modifications, both naturally occurring and artificial, are available to improve ASO pharmacologic properties. Most typical ASOs may be tailored by chemically modifying the 2’ oxygen to increase stability, nuclease resistance, and target affinity of the ASO. Modification of the phosphate backbone also can improve the pharmacokinetic and pharmacodynamic profile of the construct (24). Locked nucleic acids (LNAs), which are a synthetic nucleotide modification, drastically increase affinity to the target mRNA sequence (25). Finally, “gapmer” oligonucleotides, which are chimeric DNA and RNA constructs, have been FDA approved for a variety of indications (26).

To generate an efficacious and selective anti-IR-A ASO, we tested a variety of modified ASOs using strategies discussed above to identify the best possible anti-IR-A ASO. We identified one ASO that selectively knocks down IR-A mRNA in multiple tumor cell types with minimal impact on IR-B. This ASO may be delivered via lipofectamine or without vehicle delivery through a process known as gymnosis (27). Although we do not currently have the tools to selectively identify IR-A versus IR-B at the protein level, the ASO we designed and tested also produces a modest but significant reduction in total IR protein, indicating selective knockdown of isoform A. There is a significant gap in our knowledge of the respective functions of the IR isoforms due to a lack of tools to study each isoform individually. The ASO design we report here provides an inexpensive and easy-to-use method of targeting IR-A to better understand its functions in the context of cancer and cell physiology. We also present evidence that the anti-IR-A ASO has potential therapeutic value by inhibiting the oncogenic functions of IR-A in cancer cells.

## Methods and Materials

### Antisense Oligonucleotides

ASOs were designed by us and purchased from IDT. A list of relevant ASO designs is provided in Table 1, along with a summary of results obtained from each ASO. Scramble control sequences were generated from the GenScript siRNA Sequence Scrambler tool. We used Sigma-Aldrich’s OligoEvaluator tool to assess for intra-oligo secondary structure and primer dimers. We input the sequences for i7 and i12 in the software, in either RNA or DNA form, as four separate calculations; in all cases, neither i7 nor i12 oligos were predicted to form secondary structure or primer dimers.

**Table 1.**
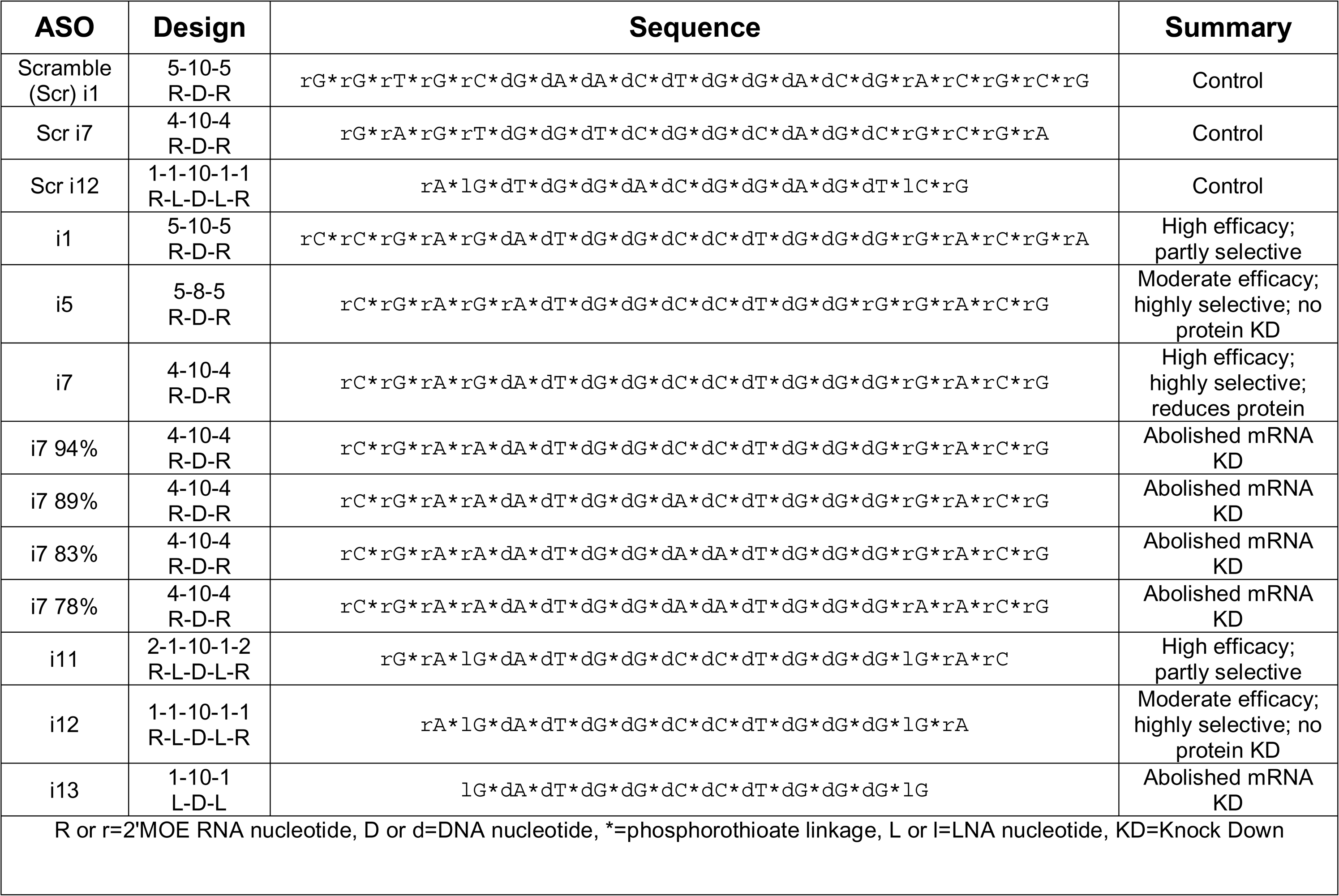
List of relevant anti-IR-A ASO variants tested, with a summary of the efficacy of each variant.

### Cell Culture, Transfection, and Gymnosis

Cell lines were obtained from ATCC (Manassas, VA), authenticated in the past year by ATCC, and tested for mycoplasma contamination twice annually. MDA-MB-231(28) (RRID:CVCL_0062) breast cancer and Hs822.T (RRID:CVCL_0933) Ewing’s sarcoma cells were maintained in DMEM (Thermo Fisher, Waltham, MA) with 10% fetal bovine serum (FBS) and 1% penicillin/streptomycin. 22Rv1(29) (RRID:CVCL_1045) cells were maintained in RPMI 1640 (ATCC) with 10% FBS and 1% penicillin/streptomycin. Media changes or passages were performed 2-3 times per week. Cells were seeded at a density of 5×10^5^ per well in 6 well plates or 1-2×10^5^ per well in 12 well plates the day prior to transfection. ASOs were transfected at the concentrations indicated in each figure via Lipofectamine RNAiMAX reagent (Thermo Fisher #13778030) per the manufacturer’s instructions. Cells were collected for lysis after 24 hours for RNA isolation or after 3 days for protein isolation unless otherwise indicated. For gymnosis experiments, cells were seeded in 12-well format. On the day after seeding, cell media was aspirated and replaced with 1 ml media containing the indicated concentration of ASO with no lipofectamine or other delivery vehicle. RNA was then isolated 24 hours later as indicated in the “RNA Isolation and Quantitative RT-PCR” section.

### RNA Isolation and Quantitative RT-PCR

Cell plates were placed on ice immediately prior to cell lysis. Cells were washed with 1x PBS, then incubated in RLT lysis buffer from Qiagen (Hilden, Germany) RNeasy Mini Kit (#74106) for 5 minutes on ice. Cells were then scraped and collected, and RNA isolation was performed via Qiagen instructions. RNA concentration and quality were assessed via the NanoDrop ND-1000 spectrophotometer (Thermo Fisher). 300 or 500 ng of RNA were used for cDNA synthesis via the iScript cDNA Synthesis Kit (BioRad, Hercules, CA #1708891). cDNA was diluted 1:5 in nuclease-free water. qPCR was performed using iTaq™ Universal SYBR® Green Supermix (BioRad #1725124) and primers listed in Suppl. Table 1 via the BioRad CFX96 real-time PCR machine. IR-A and IR-B primers were used in qPCR assays to quantify each mRNA as previously described (19). Relative expression values were calculated via the Q-Gene software from BioTechniques Software Library (30) using β-tubulin as the reference gene. Final values were multiplied by 10^3^ to increase numbers above decimal fractions.

### Protein Isolation, SDS-PAGE and Western Blot

Cell plates were placed immediately on ice prior to lysis 3 days post-transfection unless otherwise indicated, and cells were washed with 1x PBS. Cells were then incubated for 5 minutes in Pierce RIPA Lysis and Extraction Buffer (Thermo Fisher #89901) supplemented with Halt™ Protease Inhibitor Cocktail (Thermo Fisher #1861281) before scraping and collection. Protein concentration was measured via the BioRad DC Protein Assay Reagent A (#5000113) and Reagent B (#5000114) per the manufacturer’s instructions. 10-35 μg of protein was loaded with 4x Laemmli buffer (BioRad) and separated via SDS-PAGE. Primary antibodies included anti-IRβ (Cell Signaling, Danvers, MA #3025, RRID:AB_2280448), anti-β-tubulin (Cell Signaling #2146, RRID:AB_2210545), anti-pan-Akt (Cell Signaling #4691, RRID:AB_915783), anti-phospho-Akt Ser473 (Cell Signaling #4060, RRID:AB_2315049), and anti-cofilin (Cell Signaling #5175, RRID:AB_10622000). Primary antibodies were diluted 1:1000 in 5% bovine serum albumin/TBS-T and incubated at either 4^°^C overnight or room temperature for 1 hour. Horseradish peroxidase-labeled secondary antibodies (Gt anti-Rb, Jackson ImmunoResearch, West Grove, PA, #111-035-003) were diluted 1:5000 in 5% milk/TBS-T and incubated with the membrane for 1 hour at room temperature. Band intensities were measured in ImageJ after background subtraction.

### xCELLigence Cell Number Assay

Two days post-transfection, cells were detached via accutase (Sigma-Aldrich, Burlington, MA), collected in complete growth medium, and seeded into xCELLigence E-plates (Agilent, Santa Clara, CA #300600890) in technical quadruplicate at a density of 3×10^4^ cells per well. Plates were loaded into the xCELLigence RTCA DP analyzer (Agilent) and measurements were taken every 15 minutes over the time course indicated.

### BLAST analysis

To assess for off-target effects of i7 and i12 ASOs, we used the NIH Nucleotide BLAST sequence alignment program to search for sequence complementarity to human mRNA transcripts. We input FASTA sequences “CGTCCCCAGGCCATCTCG” for i7 and “TCCCCAGGCCATCT” for i12, which represent the target mRNA sequence for each. We chose the search set “Genomic + transcript databases” and selected “Human genome plus transcript (Human G+T)” from the dropdown menu. We then optimized the program selection for “Highly similar sequences” and searched, keeping all other settings as default. We used “Query Cover” as our criteria for similarity to the target transcript, as it is a readout of contiguous base pairing.

### Statistics and Rigor

Biological replicates were generated from cell cultures from separate thaws of stocks or from experiments run on separate days. Experiments comparing 2 groups were assessed for significance via two-tailed t-test, and experiments comparing more than 2 groups were analyzed using a one-way ANOVA and post-hoc Tukey’s or Dunnett’s multiple comparisons test. Alpha was set to .05 for all experiments. Error bars were generated using the mean and standard error calculated from each analysis.

## Results

### ASO Design

FDA-approved ASOs for various indications commonly feature a 5-10-5 RNA-DNA-RNA gapmer design with 2’oxygen modifications and phosphate backbone modifications (24). As such, we first tested a 5-10-5 gapmer with 2’-methoxyethyl and phosphorothioate linkage modifications, referred to as “i1”. We designed the ASO to target the exon 10-12 junction symmetrically, and this resulted in a potent and moderately selective knockdown of IR-A versus IR-B via RNAiMAX Lipofectamine transfection (Fig. 1A). IR-A mRNA levels were significantly reduced in MDA-MB-231 cells transfected with i1 as assessed by RT-qPCR, indicating that the ASO successfully induced RNase H-mediated cleavage of the target mRNA. In a “microwalk” experiment, we generated 6 variants of this ASO targeting the exon 10-12 junction site; the variants were complementary to sequences 1, 2, and 3 nucleotides both upstream and downstream of the original, symmetric i1 target site (Suppl. Table 1). We found that targeting the junction site symmetrically produced the most robust and selective knockdown of IR-A in MDA-MB-231 cells (Suppl. Fig. 1).

Substitution of 3 RNA nucleotides in each flank of the 5-10-5 gapmer resulted in nonselective knockdown of both IR-A and IR-B (data not shown). In a similar manner, longer oligonucleotides with additional nucleotides in the DNA gap or RNA wing of the ASO produced nonselective knockdowns of both IR-A and IR-B (Suppl. Fig. 2). However, truncation of the original 5-10-5 gapmer to a 4-10-4 (i7) or 5-8-5 (i5) design greatly reduced off-target IR-B mRNA reduction (Fig. 1A). In a preliminary study, we found that the 4-10-4 design, but not the 5-8-5 design, knocked down total IR protein (data not shown). We continued to characterize i7 to determine its activity against IR-A, and tested ASO variants with single LNA substitutions. We found that a variant with one LNA and one RNA in the gapmer flank, i12, had similar efficacy and selectivity to i7 (Suppl. Fig. 3). Although i1 and i11 showed a high degree of IR-A:IR-B knockdown, they induced a greater off-target knockdown of IR-B than i7 and i12. We were particularly interested in characterizing i12, since ASOs with 16 nucleotides or fewer may undergo gymnotic uptake (31).

### Dose-Response Curves

In a dose-response study of i7 and i12 versus their scramble-sequence controls, i7 and i12 significantly reduced IR-A mRNA and the IR-A:IR-B mRNA ratio up to 1 µM with minimal effect on IR-B mRNA (Fig. 1B-C). However, i7 was significantly more efficacious than i12 in reducing the IR-A:IR-B mRNA ratio.

Although i12 appeared to be less efficacious than i7, we were curious to see if its short sequence conferred greater deliverability via gymnotic uptake. When MDA-MB-231 cells were incubated with 100 nM of i7 or i12 in the absence of lipofectamine, there was no change in IR-A or IR-B mRNA levels (Suppl. Fig. 4). However, at a concentration of 5 µM, both i7 and i12 significantly reduced IR-A and IR-A:IR-B mRNA levels without affecting IR-B mRNA (Fig 2A). In this experiment, i7 reduced IR-A mRNA significantly more than i12. This was surprising, as i7 is a larger oligonucleotide, and gymnosis has been reported to occur during delivery of shorter, LNA-modified ASOs of 16 nucleotides or fewer (31). These results suggest that i7 is the most suitable candidate for IR-A knockdown, since it has greater efficacy than i12 while still being highly selective and deliverable.

### Further ASO Characterization

To test if the IR-A ASOs could tolerate any mismatched base pairing to target mRNA, we generated 4 variants of i7 with either 1, 2, 3, or 4 nucleotide substitutions. G or C nucleotides were replaced with A nucleotides throughout the ASO. We found that i7 could not tolerate a single mismatch in base pairing (Suppl. Fig. 5). This indicates that it will be important in future experiments using i7 to check for target sequence conservation in test samples to ensure ASO efficacy.

We also tested if the IR-A ASOs were efficacious against IR-A in other cancers. We found that 100 nM of either i7 or i12 reduced the IR-A:IR-B ratio with no IR-B mRNA knockdown in Hs822.T Ewing sarcoma and 22Rv1 prostate carcinoma cells (Fig. 2B, C). i7, but not i12, significantly reduced Hs822.T and 22Rv1 IR-A mRNA. The discrepancy between the significant reduction in the IR-A:IR-B mRNA ratio and no significant reduction in IR-A mRNA with i12 may be attributable to variability of IR isoform expression between replicates, as well as the lower efficacy of i12 versus i7. It should be noted that dosing for i7 and i12 was not optimized for other cell lines, and as a result, other cancer types may require higher or lower levels of IR-A ASO depending on their expression profile.

To assess for potential off-target effects of i7 other than IR-B knockdown, we used BLAST to search for sequences that are “highly similar” to the target mRNA sequence of human IR-A. Since even one nucleotide mismatch abolished i7-mediated IR-A mRNA knockdown (Suppl. Fig. 5), we filtered out results with a query coverage of less than 100%. There were no mRNA transcripts that completely matched the target mRNA sequence of i7 other than IR-A. To find the next closest related gene, we removed our query coverage filter. We found that the closest related human mRNA transcripts, including CD86, ARHGAP33, TSPAN15, and OR10P1, have 15 out of 18 contiguous matching base pairs to i7 and are potential, albeit unlikely, unintended targets. I12 has a high potential for off-target effects, as it bears 100% sequence complementarity to portions of TSPAN15, AP1M1, OR10P1, and CD86 mRNA.

### Protein Knockdown and Biological Activity

As mentioned previously, there are no antibodies available to detect IR-A specifically, and there is no established method to detect IR-A protein. However, to confirm that i7 and i12 could reduce total IR protein, we generated a dose-response curve up to 1 µM for each using lipofectamine transfection. Total IR relative to β-tubulin was determined by Western blotting at 3 days (Fig. 3A-B; Suppl. Fig. 6). At 1 µM, i7, but not i12, produced a significant reduction in total IR. Although i7 produced a significant reduction in IR-A mRNA at 100 nM, this did not result in a reduction in total IR protein at the 3-day timepoint. This finding suggests that i7 may reduce IR-A protein via translational blockage at high doses. Alternatively, a longer incubation time may be necessary for mRNA knockdown to be reflected in total IR protein levels due to the longevity of the IR protein. To test the latter, we transfected MDA-MB-231 cells with 100 nM of i7 and i12 and incubated the cells without media changes for 1 week (Fig. 3C-D). In cells transfected for 1 week with each ASO, IR-A mRNA knockdown was maintained by i7, but not i12, versus Scr i7 (Fig. 3C). A 1-week incubation with the lower concentration of i7 resulted in a greater reduction of total IR protein than the 3-day timepoint, but the decrease was not significant (p=.0682 for Scr i7 vs. i7). Thus, higher doses appear necessary for IR-A protein knockdown, and the combination of a 1 µM dose plus longer incubation time may improve protein reduction.

**Figure 3.**
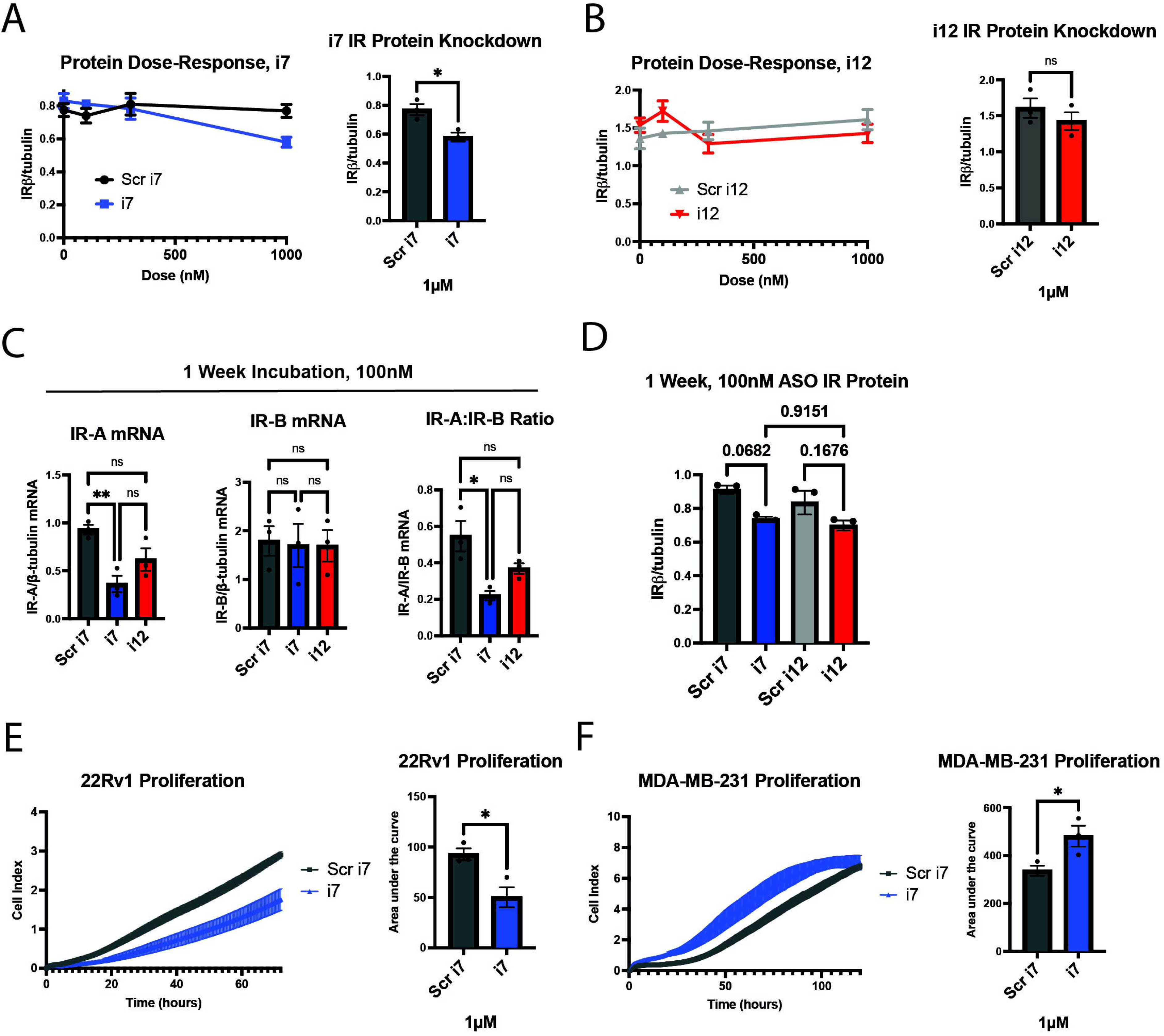
i7 ASO reduces total IRβ protein at 1 µM in MDA-MB-231 cells and alters proliferation rates in replicating cells. (A-B) A dose-response study of (A) i7 and (B) i12 with subsequent western blot analysis, showing total IR protein (detected by IRβ subunit antibody) normalized to β-tubulin. Cells were harvested 3 days post-transfection. 100 nM of either ASO did not reduce total IRβ protein, but 1 µM of i7 reduced total IRβ. (C-D) MDA-MB-231 cells were transfected with 100 nM of i7 or i12 and incubated for 1 week prior to harvest and (C) qPCR and (D) western blot analyses to assess the effect of a longer incubation time on IR isoform mRNA knockdown and total IR protein. (E-F) xCELLigence proliferation assay in which 3×10^4^ (E) 22Rv1 or (F) MDA-MB-231 cells were seeded per well of an xCELLigence E-plate and incubated for (E) 72 hours or (F) 120 hours to assess cell counts upon IR-A knockdown, shown graphically (left) and quantified via area under the curve (right).

In addition, loss of IR-A protein may be masked by the presence of baseline or upregulated IR-B protein as measured by western blot. In 22Rv1 prostate cancer cells, 1 µM i7 selectively reduced IR-A mRNA with no effect on IR-B mRNA (Suppl. Fig. 7). i7 did not reduce total IR protein as assessed by our methods; however, the main effector of IR signaling, phospho-Akt, was reduced (Suppl. Fig. 8). These results indicate that our ASO reduces IR-A protein, but this reduction may be undetectable via western blotting by our current methods.

Finally, to test for biological activity, we assessed proliferation upon IR-A knockdown since IR-A is recognized for its proliferative and tumorigenic effects in some cell types, assessed experimentally in fibroblasts, pancreatic beta cells, vascular smooth muscle cells, and endometrial carcinoma (32–36). 22Rv1 cells actively proliferated over the course of 72 hours, and IR-A knockdown with 1 µM of lipofectamine-transfected i7 reduced the cell index area under the curve by approximately 50%, indicating a robust inhibition of proliferation (Fig. 3E). Unexpectedly, IR-A knockdown via i7 increased MDA-MB-231 proliferation over the course of 120 hours (Fig. 3F). Thus, our data support the conclusion that i7 is a suitable candidate for selective knockdown of IR-A with differential effects on cell proliferation depending on the cell type. It is also possible and will be of interest in future studies to determine if IR-A has additional functions in tumor cell phenotypes such as migration or invasion properties.

## Discussion

Overexpression of IR-A confers a greater capacity for cell replication and is commonly upregulated in tumors from various tissue origins including breast, prostate, and the endometrium as outlined in Table 1 from the review by Vella and colleagues (9). In adult tissues, IR-B modulates various metabolic processes, whereas IR-A is associated with cell replication during embryogenesis (37). The metabolic function of IR-B limits its viability as a potential therapeutic target. Nonselective inhibition of the IR causes hyperglycemia, and inhibition of the insulin-like growth factor-1 receptor (IGF1R), which also contributes to glucose metabolism, commonly induces hyperglycemia in cancer clinical trials (2, 38, 39). IGF1R inhibitors have all failed in cancer clinical trials, and IR-A function has been proposed as a potential compensatory mechanism for loss of IGF1R signaling (40). Taken together, these findings indicate that IR-A is a promising anticancer target; however, its homology to IR-B renders it difficult to inhibit selectively.

No ASOs are currently approved by the FDA for cancer treatment, and drug delivery poses a major obstacle to ASO drug development due to limited uptake into target tissue (41). To circumvent this, liposomal vehicles have been developed to improve ASO delivery to tumor tissue. As Gagliardi and Ashizawa describe, one liposome-encapsulated ASO, BP1001, is currently in a Phase II clinical trial for the treatment of acute myeloid leukemia, supporting the translatability of ASOs to the clinic(41). SSOs that modulate the IR-A:IR-B mRNA ratio are a significant recent advancement in IR research (17, 42); however, reducing IR pre-mRNA splicing to IR-A results in an increase in IR-B mRNA, which correlates with worse patient outcomes in breast cancer (3). To this end, we have developed an ASO that reduces IR-A mRNA levels with little impact on IR-B expression. We hypothesize that this ASO may serve as a useful research tool to study the function of IR-A without reducing IR-B expression. Despite its effects on IR-B expression, the anti-IR-A SSO developed by Khurshid and colleagues has been successful in reducing rhabdomyosarcoma and osteosarcoma aggressiveness *in vitro* and attenuating neoangiogenesis in a mouse xenograft model of rhabdomyosarcoma (17, 42). Thus, we predict that the i7 anti-IR-A ASO will have similar or better therapeutic potential in some cancers.

Based on our BLAST analysis, i7 and especially i12 have potential off-target effects. I12’s target sequence shares 100% similarity to sequences in TSPAN15, AP1M1, OR10P1, and CD86 mRNA. This further indicates that i12 is a poor candidate for further testing as an anti-IR-A ASO. I7 bears 83% complementarity to sequences in CD86, ARHGAP33, TSPAN15, and OR10P1. Although this may be a concern, we have shown that even a single mismatch in base pairing to target IR-A mRNA results in ablation of IR-A target transcript silencing (Suppl. Fig. 5). As a result, it is unlikely that i7 will produce off-target knockdown of these mRNAs at our tested doses. In addition, sequence complementarity is not the only factor in binding affinity of an ASO to its target. mRNA secondary structure can potentially mask a target site from an RNAi agent, including ASOs and siRNAs (43). As such, target sequence similarity to other transcripts does not guarantee that there will be an off-target effect. Further studies will need to be done to confirm that i7 does not reduce mRNA levels of these other genes.

Finally, our knowledge of the IR isoforms and their function in cancer is incomplete. For instance, an emerging area of study is the role of nuclear IR and its effects on transcription (44). The IR has been found to associate with RNA polymerase II in liver cells (45); however, a potential function of nuclear IR in cancer has yet to be determined. More nebulous is the role of each IR isoform in nuclear localization and transcriptional regulation. Our proliferation studies show differential effects of IR-A knockdown in breast versus prostate cancer cell lines, and the reason for this is currently unknown. In 22Rv1 cells, IR-A knockdown reduced total levels of phospho-Akt, which is known to regulate effectors of many processes including cell division(46). This likely explains the antiproliferative effect of the IR-A ASO in this cell line; however, IR-A knockdown unexpectedly increased proliferation in MDA-MB-231 triple-negative breast cancer cells. We anticipated that, due to the known mitogenic effects of IR-A, i7 would reduce proliferation in all tested cell lines expressing the receptor. Although this result is surprising, it may be explainable by unopposed IGF1R function. IR and IGF1R heterodimerize to form hybrid receptors, and deletion of IGF1R has been shown to increase insulin sensitivity of endothelial cells (47, 48). In turn, IR knockdown may increase IGF1R homodimers, which are known to have prominent proliferative effects (49). This may also depend on levels of IGF1R expression in each cell type. The complexity of the insulin/IGF axis necessitates tools to study its components, such as IR-A. As such, our ASO will provide a means to remove IR-A from this axis and study IR-B or IGF1R in isolation. In addition, since IR-A and IGF1R have overlapping and complementary downstream effects in cancer, co-targeting both has been proposed as a potential therapeutic strategy (40). Targeting both will likely produce an overall inhibitory effect on proliferation. As such, our i7 anti-IR-A ASO may be used as a tool to selectively reduce IR-A protein and determine the effects on IR localization, transcriptional regulation, and other cell functions. It is also a candidate adjuvant drug for co-treatment of aggressive cancers.

## Acknowledgements

We extend our sincere gratitude to everyone involved in making this project possible. We thank the Rutgers Tech Transfer office and associates, including Alex Turo, Anthony Brandt, Manisha Bajpai, and others for their invaluable support in patenting the ASOs and securing funding through Rutgers TechAdvance. We thank Marie L. Mather for her expertise and support on statistical analyses conducted throughout the project. We thank Quan Shang for her essential role in managing daily activities in the lab. Finally, we thank Luis F. Almansa for chemistry expertise and consultation on this project. We also acknowledge the use of ChatGPT-4o mini, which we used only for modifying the title of this article.

## Funding Statement

This project was funded by Rutgers TechAdvance Award, ID: TA2023-0026 to TLW and New Jersey Commission on Cancer Research (NJCCR) Bridge Grant COCR23RBG003 to TLW.

## Conflict of Interest

Rutgers University has filed PCT patent application PCT/ US2024/061144 “COMPOSITIONS AND METHODS FOR TARGETING INSULIN RECEPTOR ISOFORM A (IRA)” on behalf of inventors CAG and TLW.

## Author Contributions

Christopher A. Galifi generated all data and analyses, wrote the initial text draft of the manuscript, designed the original ASO i1 and tested all variants thereafter. Ryan J. Dikdan provided invaluable consultations on the ASO design that resulted in the generation and testing of the highly selective i7 and i12 variants. Divyangi Kantak provided consultation on early oligo designs for this project. Joseph J. Bulatowicz and Krystopher Maingrette made essential intellectual contributions regarding experimental design and technique throughout the project. Samuel I. Gunderson provided essential expert consultation on ASO design and testing throughout the duration of the project and guided the ASO microwalk and gymnosis experiments. Teresa L. Wood guided the entirety of this project from its conception and co-invented the original ASO concept and design. Both SIG and TLW read and edited the manuscript.

## Nomenclature

ASO: antisense oligonucleotide
BSA: bovine serum albumin
FBS: fetal bovine serum
IGF1R: insulin-like growth factor 1 receptor
IGF2: insulin-like growth factor 2
IR: insulin receptor
LNA: locked nucleic acid
SSO: splice-switching oligonucleotide

## Data Availability Statement

The datasets generated for this study will be deposited on OSF and the link will be made available upon acceptance and publication.

## References

1. Kahn CR, White MF. The insulin receptor and the molecular mechanism of insulin action. J Clin Invest. 1988;82(4):1151–6.

2. Galifi CA, Wood TL. Insulin-like growth factor-1 receptor crosstalk with integrins, cadherins, and the tumor microenvironment: sticking points in understanding IGF1R function in cancer. Endocr Relat Cancer. 2023;30(10).

3. Gradishar WJ, Yardley DA, Layman R, Sparano JA, Chuang E, Northfelt DW, et al. Clinical and Translational Results of a Phase II, Randomized Trial of an Anti-IGF-1R (Cixutumumab) in Women with Breast Cancer That Progressed on Endocrine Therapy. Clin Cancer Res. 2016;22(2):301–9.

4. Cao J, Yee D. Disrupting Insulin and IGF Receptor Function in Cancer. Int J Mol Sci. 2021;22(2).

5. Obeagu EI, Obeagu GU. Breast cancer: A review of risk factors and diagnosis. Medicine (Baltimore). 2024;103(3):e36905.

6. Centers for Disease Control and Prevention. U.S. Cancer Statistics Female Breast Cancer Stat Bite 2024.

7. Frasca F, Pandini G, Scalia P, Sciacca L, Mineo R, Costantino A, et al. Insulin receptor isoform A, a newly recognized, high-affinity insulin-like growth factor II receptor in fetal and cancer cells. Mol Cell Biol. 1999;19(5):3278–88.

8. Belfiore A, Malaguarnera R, Vella V, Lawrence MC, Sciacca L, Frasca F, et al. Insulin Receptor Isoforms in Physiology and Disease: An Updated View. Endocr Rev. 2017;38(5):379–431.

9. Vella V, Milluzzo A, Scalisi NM, Vigneri P, Sciacca L. Insulin Receptor Isoforms in Cancer. Int J Mol Sci. 2018;19(11).

10. An W, Hall C, Li J, Hung A, Wu J, Park J, et al. Activation of the insulin receptor by insulin-like growth factor 2. Nat Commun. 2024;15(1):2609.

11. Belfiore A, Malaguarnera R. Insulin receptor and cancer. Endocr Relat Cancer. 2011;18(4):R125–47.

12. Aljada A, Saleh AM, Al-Aqeel SM, Shamsa HB, Al-Bawab A, Al Dubayee M, et al. Quantification of insulin receptor mRNA splice variants as a diagnostic tumor marker in breast cancer. Cancer Biomark. 2015;15(5):653–61.

13. Huang J, Morehouse C, Streicher K, Higgs BW, Gao J, Czapiga M, et al. Altered expression of insulin receptor isoforms in breast cancer. PLoS One. 2011;6(10):e26177.

14. Harrington SC, Weroha SJ, Reynolds C, Suman VJ, Lingle WL, Haluska P. Quantifying insulin receptor isoform expression in FFPE breast tumors. Growth Horm IGF Res. 2012;22(3-4):108–15.

15. Vella V, Giuliano M, La Ferlita A, Pellegrino M, Gaudenzi G, Alaimo S, et al. Novel Mechanisms of Tumor Promotion by the Insulin Receptor Isoform A in Triple-Negative Breast Cancer Cells. Cells. 2021;10(11).

16. Zhang H, Fagan DH, Zeng X, Freeman KT, Sachdev D, Yee D. Inhibition of cancer cell proliferation and metastasis by insulin receptor downregulation. Oncogene. 2010;29(17):2517–27.

17. Khurshid S, Montes M, Comiskey DF, Jr., Shane B, Matsa E, Jung F, et al. Splice-switching of the insulin receptor pre-mRNA alleviates tumorigenic hallmarks in rhabdomyosarcoma. NPJ Precis Oncol. 2022;6(1):1.

18. Havens MA, Hastings ML. Splice-switching antisense oligonucleotides as therapeutic drugs. Nucleic Acids Res. 2016;44(14):6549–63.

19. Flannery CA, Rowzee AM, Choe GH, Saleh FL, Radford CC, Taylor HS, et al. Development of a Quantitative PCR Assay for Detection of Human Insulin-Like Growth Factor Receptor and Insulin Receptor Isoforms. Endocrinology. 2016;157(4):1702–8.

20. Rowzee AM, Ludwig DL, Wood TL. Insulin-like growth factor type 1 receptor and insulin receptor isoform expression and signaling in mammary epithelial cells. Endocrinology. 2009;150(8):3611–9.

21. Perks CM, Zielinska HA, Wang J, Jarrett C, Frankow A, Ladomery MR, et al. Insulin Receptor Isoform Variations in Prostate Cancer Cells. Front Endocrinol (Lausanne). 2016;7:132.

22. Dias N, Stein CA. Antisense oligonucleotides: basic concepts and mechanisms. Mol Cancer Ther. 2002;1(5):347–55.

23. Crooke ST. Molecular Mechanisms of Antisense Oligonucleotides. Nucleic Acid Ther. 2017;27(2):70–7.

24. Egli M, Manoharan M. Chemistry, structure and function of approved oligonucleotide therapeutics. Nucleic Acids Res. 2023;51(6):2529–73.

25. Braasch DA, Corey DR. Locked nucleic acid (LNA): fine-tuning the recognition of DNA and RNA. Chem Biol. 2001;8(1):1–7.

26. Chan L, Yokota T. Development and Clinical Applications of Antisense Oligonucleotide Gapmers. Methods Mol Biol. 2020;2176:21–47.

27. Stein CA, Hansen JB, Lai J, Wu S, Voskresenskiy A, Hog A, et al. Efficient gene silencing by delivery of locked nucleic acid antisense oligonucleotides, unassisted by transfection reagents. Nucleic Acids Res. 2010;38(1):e3.

28. Cailleau R, Young R, Olive M, Reeves WJ, Jr. Breast tumor cell lines from pleural effusions. J Natl Cancer Inst. 1974;53(3):661–74.

29. Sramkoski RM, Pretlow TG, 2nd, Giaconia JM, Pretlow TP, Schwartz S, Sy MS, et al. A new human prostate carcinoma cell line, 22Rv1. In Vitro Cell Dev Biol Anim. 1999;35(7):403–9.

30. Muller PY, Janovjak H, Miserez AR, Dobbie Z. Processing of gene expression data generated by quantitative real-time RT-PCR. Biotechniques. 2002;32(6):1372–4, 6, 8-9.

31. Soifer HS, Koch T, Lai J, Hansen B, Hoeg A, Oerum H, et al. Silencing of gene expression by gymnotic delivery of antisense oligonucleotides. Methods Mol Biol. 2012;815:333–46.

32. Massimino M, Sciacca L, Parrinello NL, Scalisi NM, Belfiore A, Vigneri R, et al. Insulin Receptor Isoforms Differently Regulate Cell Proliferation and Apoptosis in the Ligand-Occupied and Unoccupied State. Int J Mol Sci. 2021;22(16).

33. Wang C, Su K, Zhang Y, Zhang W, Zhao Q, Chu D, et al. IR-A/IGF-1R-mediated signals promote epithelial-mesenchymal transition of endometrial carcinoma cells by activating PI3K/AKT and ERK pathways. Cancer Biol Ther. 2019;20(3):295–306.

34. Morrione A, Valentinis B, Xu SQ, Yumet G, Louvi A, Efstratiadis A, et al. Insulin-like growth factor II stimulates cell proliferation through the insulin receptor. Proc Natl Acad Sci U S A. 1997;94(8):3777–82.

35. Escribano O, Gomez-Hernandez A, Diaz-Castroverde S, Nevado C, Garcia G, Otero YF, et al. Insulin receptor isoform A confers a higher proliferative capability to pancreatic beta cells enabling glucose availability and IGF-I signaling. Mol Cell Endocrinol. 2015;409:82–91.

36. Gomez-Hernandez A, Escribano O, Perdomo L, Otero YF, Garcia-Gomez G, Fernandez S, et al. Implication of insulin receptor A isoform and IRA/IGF-IR hybrid receptors in the aortic vascular smooth muscle cell proliferation: role of TNF-alpha and IGF-II. Endocrinology. 2013;154(7):2352–64.

37. Andres SF, Simmons JG, Mah AT, Santoro MA, Van Landeghem L, Lund PK. Insulin receptor isoform switching in intestinal stem cells, progenitors, differentiated lineages and tumors: evidence that IR-B limits proliferation. J Cell Sci. 2013;126(Pt 24):5645–56.

38. Maddux BA, Chang YN, Accili D, McGuinness OP, Youngren JF, Goldfine ID. Overexpression of the insulin receptor inhibitor PC-1/ENPP1 induces insulin resistance and hyperglycemia. Am J Physiol Endocrinol Metab. 2006;290(4):E746–9.

39. Kasprzak A. Insulin-Like Growth Factor 1 (IGF-1) Signaling in Glucose Metabolism in Colorectal Cancer. Int J Mol Sci. 2021;22(12).

40. Buck E, Gokhale PC, Koujak S, Brown E, Eyzaguirre A, Tao N, et al. Compensatory insulin receptor (IR) activation on inhibition of insulin-like growth factor-1 receptor (IGF-1R): rationale for cotargeting IGF-1R and IR in cancer. Mol Cancer Ther. 2010;9(10):2652–64.

41. Gagliardi M, Ashizawa AT. The Challenges and Strategies of Antisense Oligonucleotide Drug Delivery. Biomedicines. 2021;9(4).

42. Khurshid S, Venkataramany AS, Montes M, Kipp JF, Roberts RD, Wein N, et al. Employing splice-switching oligonucleotides and AAVrh74.U7 snRNA to target insulin receptor splicing and cancer hallmarks in osteosarcoma. Mol Ther Oncol. 2024;32(4):200908.

43. Gredell JA, Berger AK, Walton SP. Impact of target mRNA structure on siRNA silencing efficiency: A large-scale study. Biotechnol Bioeng. 2008;100(4):744–55.

44. Batista TM, Cederquist CT, Kahn CR. The insulin receptor goes nuclear. Cell Res. 2019;29(7):509–11.

45. Hancock ML, Meyer RC, Mistry M, Khetani RS, Wagschal A, Shin T, et al. Insulin Receptor Associates with Promoters Genome-wide and Regulates Gene Expression. Cell. 2019;177(3):722–36 e22.

46. Pal I, Mandal M. PI3K and Akt as molecular targets for cancer therapy: current clinical outcomes. Acta Pharmacol Sin. 2012;33(12):1441–58.

47. Turvey SJ, McPhillie MJ, Kearney MT, Muench SP, Simmons KJ, Fishwick CWG. Recent developments in the structural characterisation of the IR and IGF1R: implications for the design of IR-IGF1R hybrid receptor modulators. RSC Med Chem. 2022;13(4):360–74.

48. Abbas A, Imrie H, Viswambharan H, Sukumar P, Rajwani A, Cubbon RM, et al. The insulin-like growth factor-1 receptor is a negative regulator of nitric oxide bioavailability and insulin sensitivity in the endothelium. Diabetes. 2011;60(8):2169–78.

49. Riedemann J, Macaulay VM. IGF1R signalling and its inhibition. Endocr Relat Cancer. 2006;13 Suppl 1:S33–43.

