## Supplemental Figures for "Identifying Antisense Oligonucleotides for Targeted Inhibition of Insulin Receptor Isoform A"

Suppl. Table 1. List of primers or oligos synthesized by IDT and utilized during testing of the anti-IR-A ASO.

| Product | Sequence (5’→3’) or Catalog # | Exon Target |
|---|---|---|
| Human IR-A Forward Primer | TTT TCG TCC CCA GGC CAT C | 10-12 |
| Human IR-B Forward Primer | CCC CAG AAA AAC CTC TTC AGG | 10-11 |
| Human IR Reverse Primer | GTC ACA TTC CCA ACA TCG CC | N/A |
| Human β-tubulin | #NM_178014 | 1-2 |
| i1 +1 | rT*rC*rC*rG*rA*dG*dA*dT*dG*dG*dC*dC*dT*dG*dG*rG*rG*rA*rC*rG | 10-12 |
| i1 +2 | rT*rT*rC*rC*rG*dA*dG*dA*dT*dG*dG*dC*dC*dT*dG*rG*rG*rG*rA*rC | 10-12 |
| i1 +3 | rT*rT*rT*rC*rC*dG*dA*dG*dA*dT*dG*dG*dC*dC*dT*rG*rG*rG*rG*rA | 10-12 |
| i1 −1 | rC*rG*rA*rG*rA*dT*dG*dG*dC*dC*dT*dG*dG*dG*dG*rA*rC*rG*rA*rA | 10-12 |
| i1 −2 | rG*rA*rG*rA*rT*dG*dG*dC*dC*dT*dG*dG*dG*dG*dA*rC*rG*rA*rA*rA | 10-12 |
| i1 −3 | rA*rG*rA*rT*rG*dG*dC*dC*dT*dG*dG*dG*dG*dA*dC*rG*rA*rA*rA*rA | 10-12 |
| i6 | rT*rC*rC*rG*rA*dG*dA*dT*dG*dG*dC*dC*dT*dG*dG*dG*dG*rA*rC*rG*rA*rA | 10-12 |
| i8 | rT*rC*rC*rG*rA*rG*dA*dT*dG*dG*dC*dC*dT*dG*dG*dG*rG*rA*rC*rG*rA*rA | 10-12 |
| For ASO oligos: r=2'MOE RNA nucleotide, d=DNA nucleotide, *=phosphorothioate linkage | | |

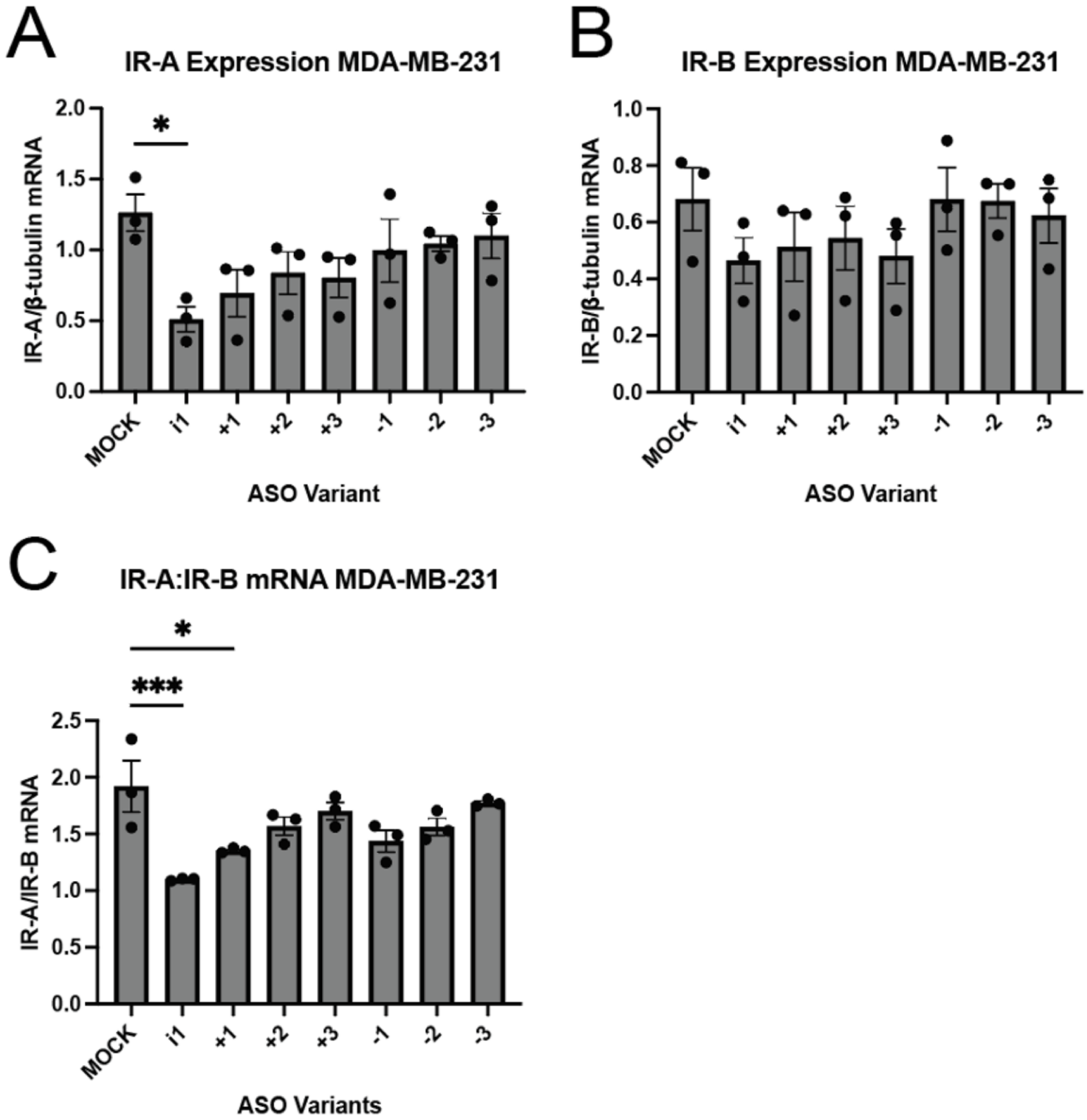


Supplemental Figure 1. ASO microwalk. Overlapping 20-nucleotide ASOs were designed in 1-nucleotide steps (+1, +2, +3, −1, −2, −3) based on the sequence of i1. Sequences are shown in Suppl. Table 1. mRNA levels of (A) IR-A, (B) IR-B, and (C) the IR-A:IR-B mRNA ratio were assessed in MDA-MB-231 cells via RT-qPCR after lipofectamine transfection as described in the methods.

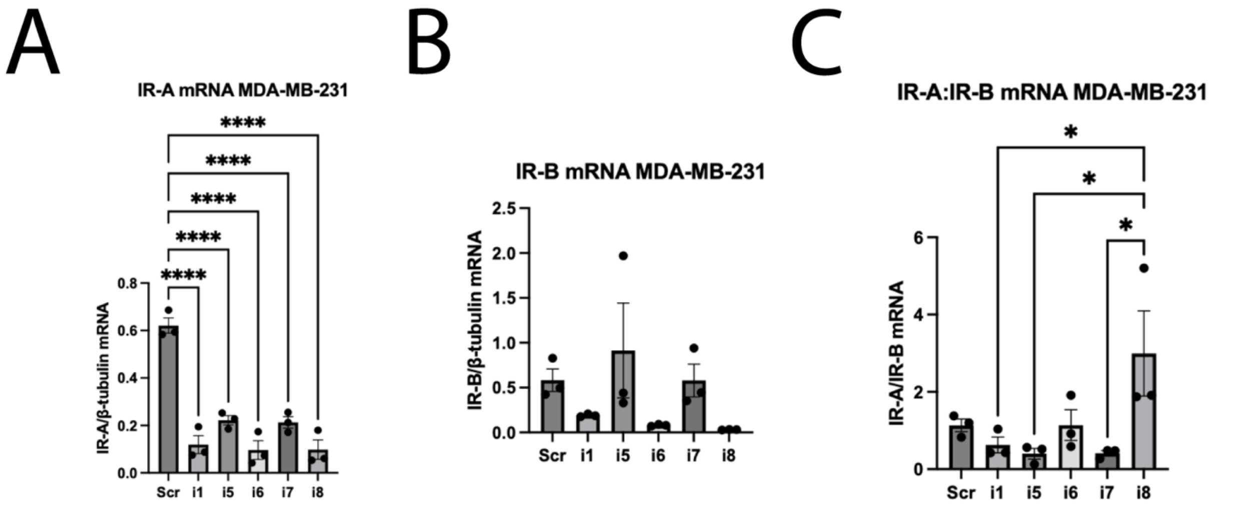


Supplemental Figure 2. Comparison of anti-IR-A oligo variants i1 and i5-i8. qPCR analysis was performed after lipofectamine transfection as described in the methods. i6 and i8 designs are listed in Suppl. Table 1. (A) IR-A, (B) IR-B, and (C) the IR-A:IR-B mRNA ratio were assessed in MDA-MB-231 cells.

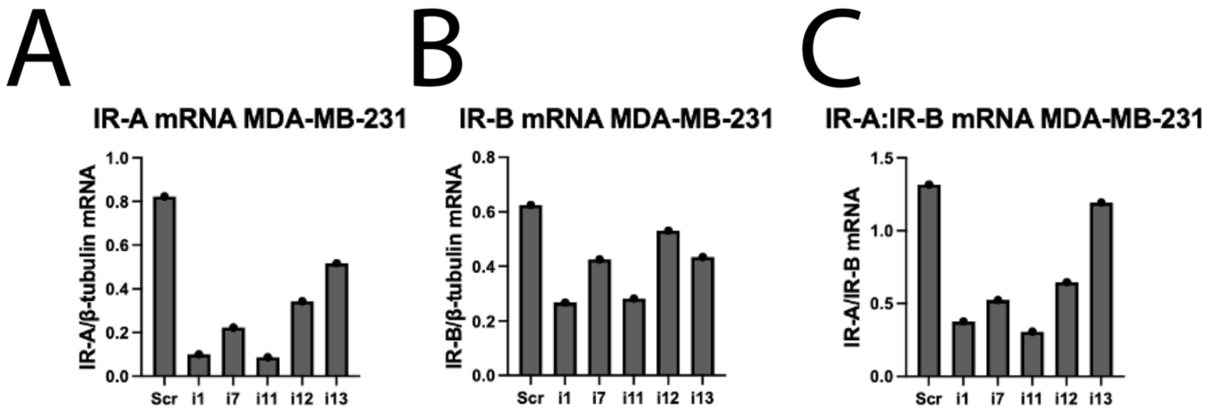


Supplemental Figure 3. Comparison of anti-IR-A oligo variants i1 and i7 to LNA-modified variants i11-i13. qPCR analysis was performed after lipofectamine transfection as described in the methods. i11-i13 designs are listed in Table 1 in the main text. (A) IR-A, (B) IR-B, and (C) the IR-A:IR-B mRNA ratio were assessed in MDA-MB-231 cells.

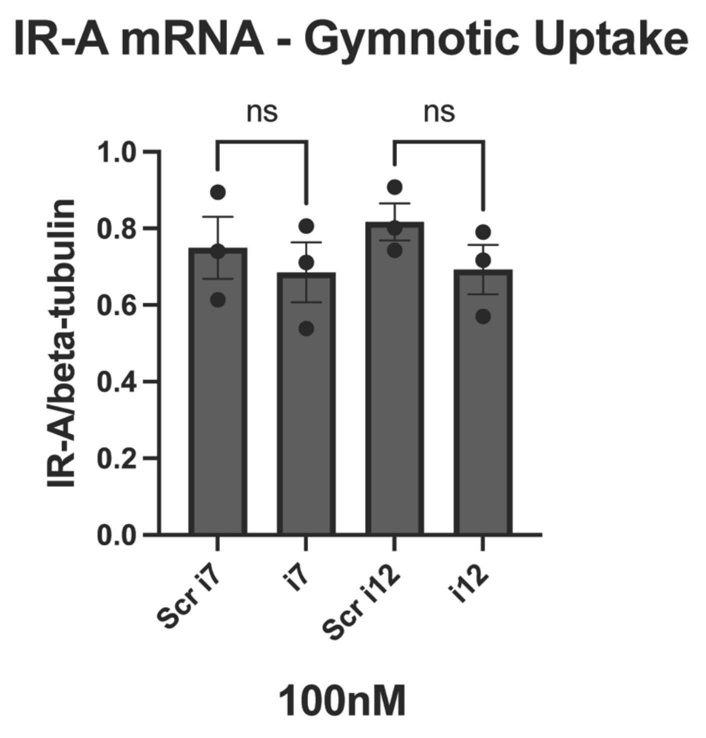


Supplemental Figure 4. Assessment of gymnotic uptake of i7 and i12 at 100 nM in MDA-MB-231 cells. Cells were incubated with each oligo as described in the methods.

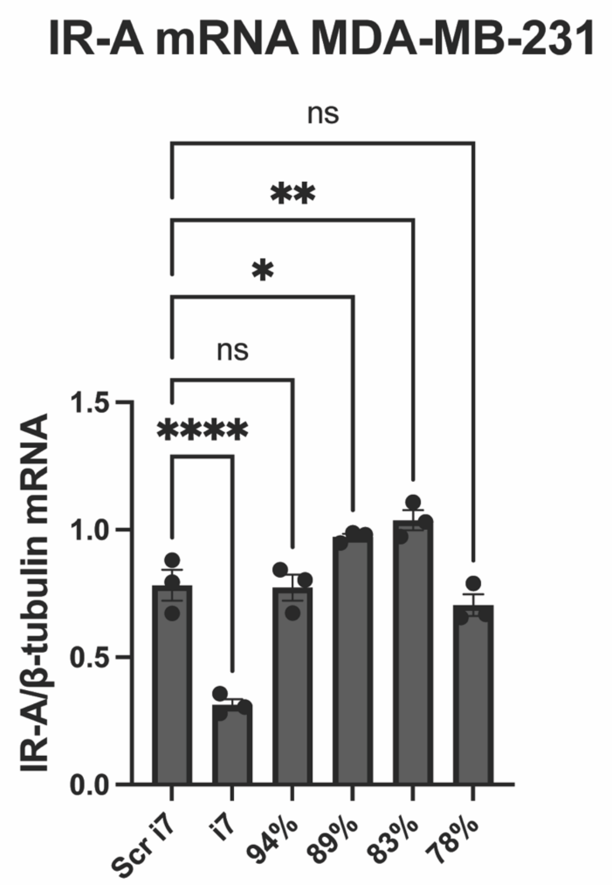


Supplemental Figure 5. Further characterization of the anti-IR-A oligos in MDA-MB-231 cells shows intolerance of target sequence mismatches for silencing efficacy. i7 oligo variants bearing 1, 2, 3 or 4 sequence mismatches, represented in the figure by their respective percent complementarity to the target human IR-A mRNA, were tested to assess for silencing efficacy at 100 nM. Oligos were transfected via RNAiMAX lipofectamine as described in the methods. Sequences for each oligo are found in Table 1 in the main text. Statistical analysis was performed via one-way ANOVA and subsequent Dunnett’s multiple comparisons test.

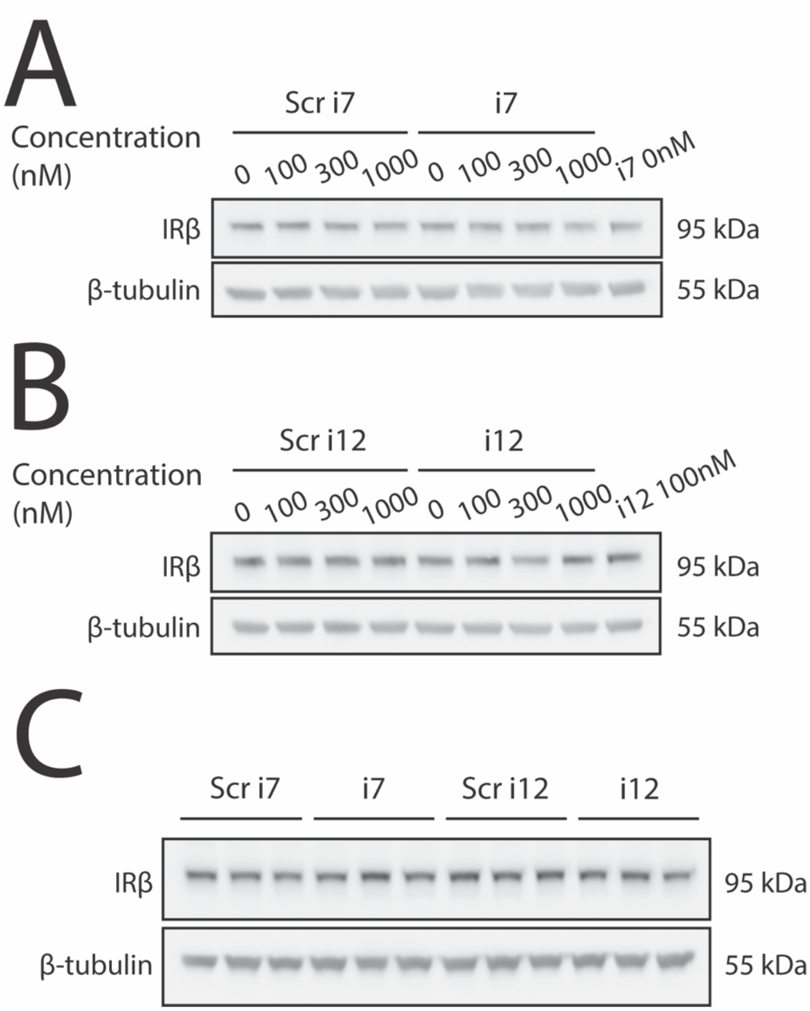


Supplemental Figure 6. Raw western blot data of i7 and i12 dose response curves. (A) i7 and (B) i12 were lipofectamine transfected into MDA-MB-231 cells as described in the methods at doses of 0, 100, 300, and 1000nM versus their scramble-sequence controls. Each replicate of these experiments was run on separate blots, and antibody concentrations and exposure times for chemiluminescence were kept consistent between replicates. The final lane of both blots was loaded with a common sample from another replicate from each experiment, so that band intensities could be compared across blots. (C) MDA-MB-231 cells were transfected with 100 nM of i7 and i12 versus scramble-sequence controls and incubated for 1 week prior to western blot analysis to determine the effect of a longer incubation time on knockdown efficacy.

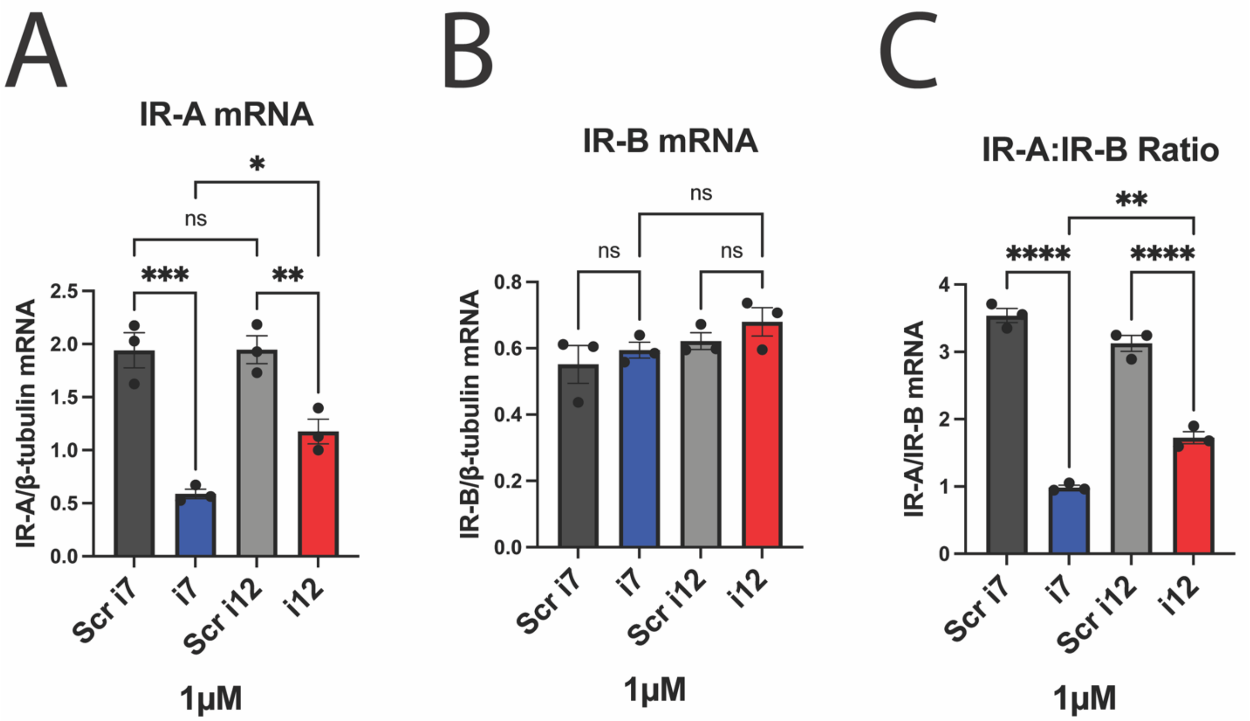


Supplemental Figure 7. 1 μM i7 selectively reduces IR-A mRNA versus IR-B post-lipofectamine transfection of 22Rv1 cells. Cells were transfected, and isolated RNA was used for RT-qPCR as described in the methods. (A) IR-A, (B) IR-B, and (C) the IR-A:IR-B mRNA ratio were assessed.

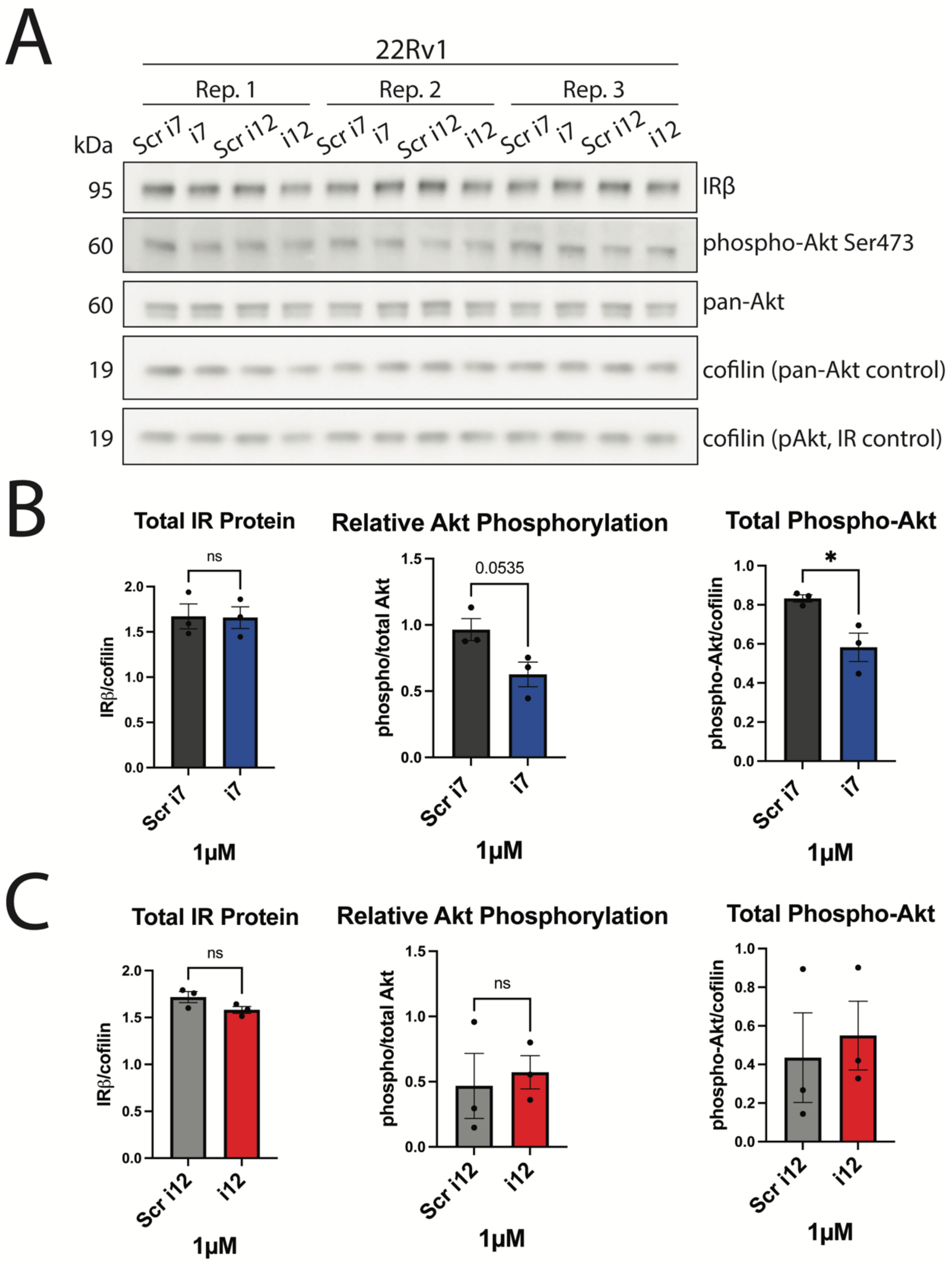


Supplemental Figure 8. 1 μM i7 reduces phospho-Akt levels in 22Rv1 cells post-lipofectamine transfection. Transfection with i7 and i12, protein isolation, and western blots were performed as described in the methods. Protein was isolated 3 days post-transfection. (A) Raw bands from Western blots of each treatment group, probed for total IR (β subunit), phospho-Akt, total Akt, and cofilin loading control protein levels were analyzed. The effects of (B) i7 and (C) i12 versus scramble-sequence controls on Akt and total IR were analyzed via two-tailed t-test.
